# PAIR-link: a ligation-free strategy that joins two RNAs into a single sequence-preserving Chimeric cDNA

**DOI:** 10.64898/2026.05.29.728768

**Authors:** Shengyi Fei, Dehao Kong, Boxuan Simen Zhao

## Abstract

Physical proximity between RNA molecules, and between the proteins that bind them, underlies much of post-transcriptional regulation, yet the methods that convert these relationships into sequence do so at a cost. Established RNA-RNA interaction maps techniques depend on crosslinking, fragmentation, and/or proximity ligation, so each partner survives only as a short chimeric read in which full identity and sequence are not preserved. We describe PAIR-link (Priming Annealed Isothermal RNA linking), a ligation-free and fragmentation-free strategy that joins two separately transcribed RNAs into a single contiguous cDNA while retaining the sequence of both partners. Two engineered RNAs carry complementary tails at their 3’ ends. These tails anneal and prime one another, and an isothermal Bst-family reverse transcriptase extends across the junction in both directions to copy both RNAs into one chimeric cDNA that can be amplified by PCR and read directly by Sanger sequencing. We first established the chemistry in vitro using synthesized oligonucleotides. Because the reaction requires a defined 3’ end, we drove the barcode RNAs from RNA polymerase III promoters, whose transcripts terminate at a defined site, rather than from polyadenylated polymerase II transcripts. We then implemented PAIR-link in human cells as a reconstituted ribonucleoprotein in which each barcode RNA is tethered to one half of a split reporter protein (split-cpHaloTag or split-TurboID), so that reconstitution of the split protein brings the two RNAs together and facilitates linking. Bead-intact pull-down preserves the complex through reverse transcription and amplification. We recovered the expected sequence-verified chimeric product for reporters directed independently to the nucleus and to the cytoplasm, showing that the method operates across distinct subcellular environments. To our knowledge, PAIR-link is the first method to fuse two separately produced RNAs into a single chimeric cDNA molecule with full sequences of RNA preserved, without crosslinking, fragmentation, or ligation. We discuss the current scope of the approach, which is limited to RNAs with defined 3’ ends, and outline routes toward future native transcripts and protein-interaction mapping.

## Introduction

The spatial organization of RNA and of RNA-binding proteins is a core feature of gene regulation. Small RNAs find their targets, snoRNAs guide modification, viral genomes dimerize, and ribonucleoprotein (RNP) complexes assemble in defined compartments, all through transient or stable physical proximity. Mapping which RNAs and which proteins are close to one another, and where in the cell that happens, is therefore central to understanding how the transcriptome is controlled.

Most transcriptome-scale methods for capturing RNA-RNA proximity share a common logic. Interacting RNAs are crosslinked, either through a bound protein (CLASH, hiCLIP) or with a small-molecule intercalator such as psoralen (PARIS, SPLASH, LIGR-seq, COMRADES), the bound or duplexed RNA is fragmented, and the two arms are joined by proximity ligation to create a chimeric molecule that is then sequenced (Helwak et al. 2013; Sugimoto et al. 2015; Lu et al. 2016; Aw et al. 2016; Sharma et al. 2016; Ramani et al. 2015; Cai et al. 2020). This design has been productive, but it carries two intrinsic limitations. First, fragmentation means the partners are recovered only as short reads, so the full-length identity and internal sequence of each RNA are not preserved in a single contiguous molecule. Second, the ligation step is inefficient and biased, and in a typical experiment only a small percentage of reads report a true interaction (Travis et al. 2014; Gabryelska et al. 2021).

Ligation-free chemistries do exist, but they were developed for a different purpose. Template-switching reverse transcriptases, used in SMART-seq and related single-cell protocols, and engineered retroelement RTs used in Ordered Two-Template Relay (OTTR), add a synthetic adapter to a cDNA end during reverse transcription without ligation (Picelli et al. 2013; Upton et al. 2021). These reactions join an RNA to a defined oligonucleotide that the experimenter supplies; they do not join two distinct, separately transcribed cellular RNAs to each other on the basis of their physical proximity. To our knowledge, no published method links two separately produced RNA molecules into a single, sequence-preserved cDNA without crosslinking, fragmentation, or ligation.

Here we introduce PAIR-link to fill that gap. Two engineered RNAs are given complementary 3’ tails so that, when the two molecules are brought into proximity, their tails anneal and a Bst-family reverse transcriptase extends across the annealed junction. The product is one continuous cDNA that carries the sequence of both RNAs, can be amplified by conventional PCR, and can be read directly by Sanger sequencing. Because the chemistry depends only on proximity of the two 3’ ends, it can be coupled to a reconstituted RNP in which each RNA is tethered to a protein. In that configuration the chimeric product reports that two proteins, and therefore the cellular events that bring them together, were close at the time of capture. We use localization tags to direct the reporter to the nucleus or to the cytoplasm and show that it produces correct, sequence-verified products in both, establishing PAIR-link as a proximity readout that functions across subcellular environments.

## Results

### Design principle of PAIR-link

PAIR-link uses two engineered RNAs, which we refer to as the pre and post barcode RNAs (Figure 1A). Each RNA carries a 5’ PCR handle, a barcode region that records its identity, and a short complementary region (denoted F1 and F1c) near its 3’ end. When the two molecules are close, the F1 and F1c regions anneal in an antiparallel orientation, so that the 3’ end of each RNA is positioned on the body of the other. In this mutual-priming configuration a Bst-family DNA polymerase, which can read either RNA or DNA as template, extends from both 3’ ends and copies both RNAs into a single double-stranded chimera. Amplification with primers to the two 5’ PCR handles yields a defined product whose sequence contains both barcodes in a fixed order, which is read directly by Sanger sequencing. No covalent ligation, RNA fragmentation, or chemical crosslinking is used at any step, so the full sequence of both partners is preserved in one contiguous molecule, in contrast to crosslink-fragment-ligate approaches.

**Figure 1.**
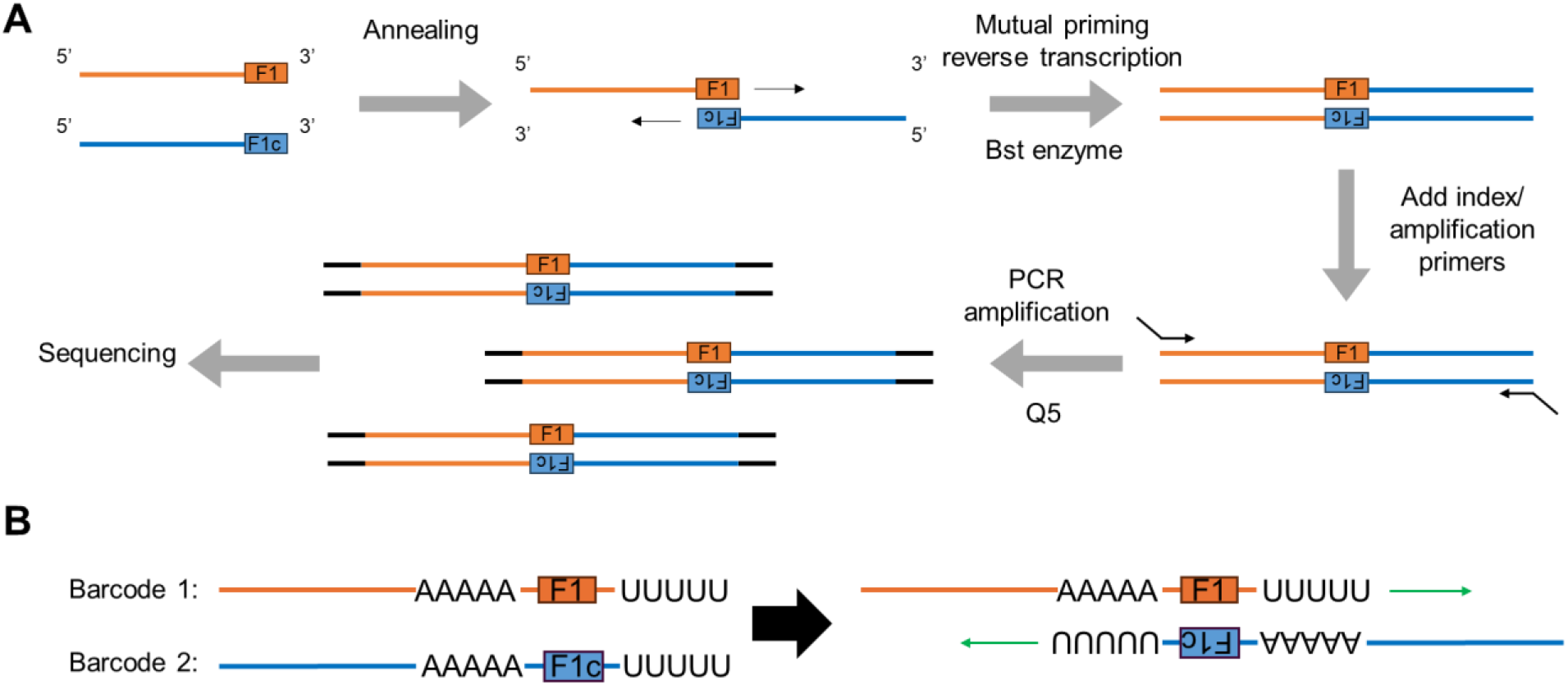
Design principle of PAIR-link. *(A)* Core linking chemistry. The two barcode RNAs each carry a 5’ PCR handle, an identity barcode, and a short complementary tag (F1, F1c) near the 3’ end. The tags anneal in antiparallel, placing each 3’ end on the body of the other RNA, and a Bst-family polymerase (which reads RNA or DNA template) extends from both ends (mutual priming) to copy both RNAs into one chimera. PCR with primers to the two 5’ handles gives a defined product containing both barcodes, read directly by Sanger sequencing. No crosslinking, fragmentation or ligation is used, and the sequence of both partners is preserved. *(B)* Barcode architecture and the defined-3’-end requirement. The annealing region sits just upstream of a defined 3’ terminus. Pol III transcripts, which terminate at a short oligo-U signal (for example hU6 is shown here), provide the required defined 3’ end, so mutual primer RT could proceed to produce complete chimeric cDNA.

Because the polymerase extends from a free 3’ end, the barcode RNA must terminate at a defined position rather than in a heterogeneous tail (Figure 1B). Polymerase II transcripts, which end in a polyadenylated tail, are therefore unsuitable, whereas RNA polymerase III transcripts, which terminate at a short run of uridines, present the defined 3’ end the reaction needs. We accordingly placed the F1 or F1c tag immediately upstream of the natural Pol III terminator and drove the barcode RNAs from the human U6 or U1 small-RNA promoters, giving transcripts that end in a defined U-rich sequence. This choice is what allows the chemistry to act on RNAs made inside the cell rather than only on synthetic oligonucleotides.

To read proximity in cells we tether each barcode RNA to one half of a split reporter protein, so that the two RNAs are brought together only when the split protein reconstitutes (Figure 1C). One RNA carries five copies of the phage P22N21 boxB hairpin and is bound by the cognate P22N peptide (Cilley and Williamson 2003), and the other carries four copies of the TAR element and is bound by a Tat peptide (Smith et al. 2000). Each peptide is fused to one fragment of a split protein, either split-cpHaloTag (Wilhelm et al. 2025) or split-TurboID (Cho et al. 2020), which also provides the handle for pull-down once reconstituted, on HaloTag ligand beads or, after biotinylation, on streptavidin beads. When the two protein fragments come together their tethered RNAs are co-concentrated, the F1 and F1c tags anneal, and linking proceeds. The complex is recovered on beads and retained through annealing, reverse transcription and the first PCR, so that proximity established in the cell is preserved during processing.

### In vitro establishment and optimization of the linking chemistry

We first asked whether two complementary barcode RNAs could be annealed and copied into a defined chimeric product (Figure 2). Synthetic barcode oligonucleotides bearing the F1 and F1c tags were annealed by controlled cooling, reverse transcribed with Bst3.0 across a temperature and time series (Figure 2A), and amplified by PCR with primers to the handles. A product of the expected size, near 96 base pairs, was obtained, and sequencing of colonies with PCR products cloned into their plasmids showed that the correct chimera formed but represented at best about half of the clones, with the remainder reflecting mispriming or incomplete extension (Figure 2B). Bst3.0 thus established feasibility but did not give a clean enough product for routine direct readout.

**Figure 2.**
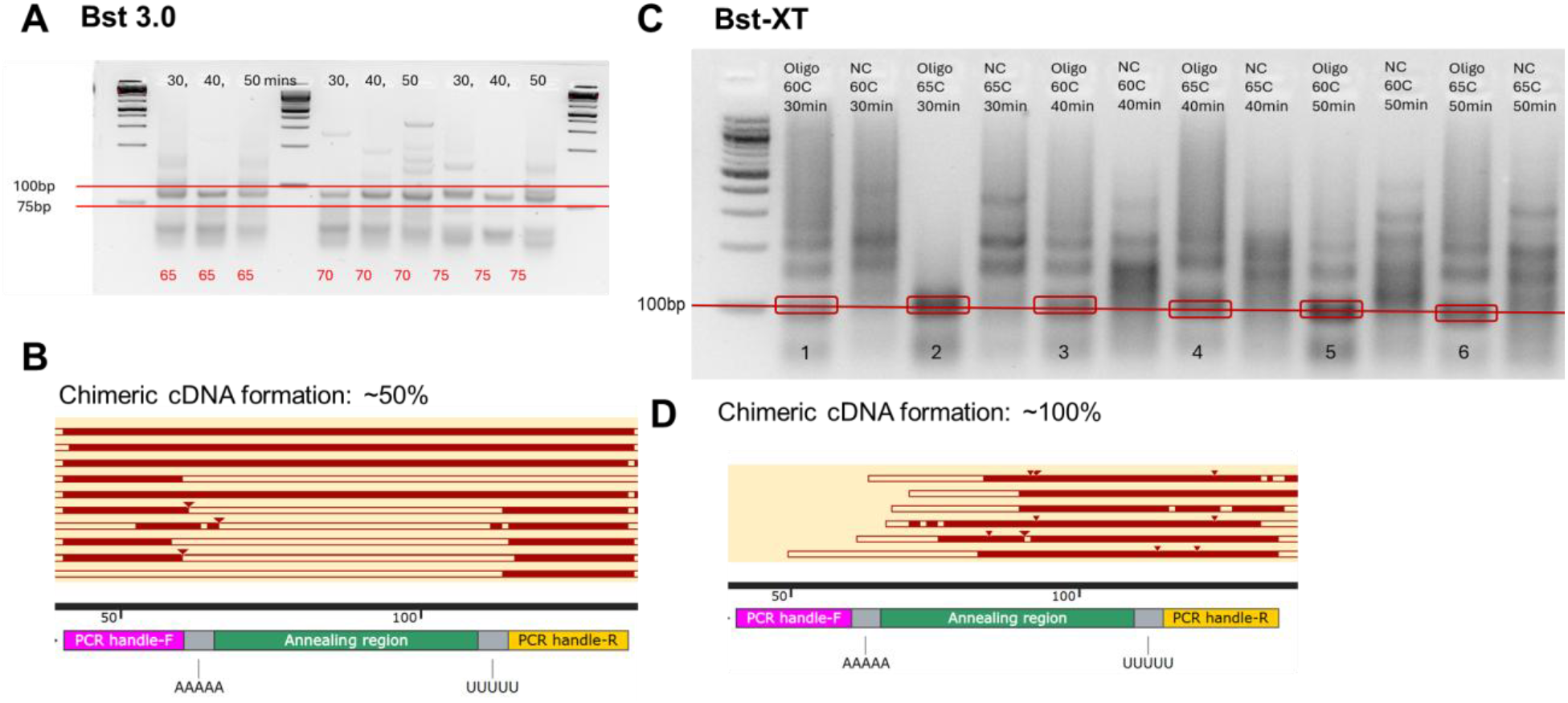
In vitro establishment and enzyme optimization. *(A)* Gel of Bst3.0 reverse transcription and PCR on synthetic barcode oligonucleotides across a temperature and time series; the target product is near 96 base pairs. *(B)* Sequence analysis of Bst3.0 products cloned into plasmids. Colony sequencing result shows correct chimera formation but accounts for about half of the molecules. *(C)* Gel of BstXT products under matched temperature and time conditions, with no-template controls; 65 C for 30 min is sufficient. *(D)* Sequence analysis of Bst-XT products cloned into plasmids. Colony sequencing result shows essentially complete and correct chimera formation.

We therefore tested a newly engineered enzyme, BstXT, under matched conditions, with no-template controls included for each temperature and time (Figure 2C). BstXT raised the fraction of correctly formed chimera to essentially complete, and 65 C for 30 min was sufficient. The amplified product could be gel-excised and Sanger sequenced directly, without cloning, and the traces confirmed the designed junction with the increased fidelity expected from the cleaner reaction (Figure 2D). We selected BstXT for all subsequent cellular work. In cells we used a longer bead-intact reaction, 65 C for 60 min, to allow for the lower effective concentration of the tethered RNAs.

### PAIR-link reads proximity of cellularly transcribed RNAs

We next applied the chemistry to barcode RNAs transcribed inside HEK293T cells and brought together by a split reporter protein (Figure 3A). Cells were transfected with the tethered barcode RNAs and the two split-protein fragments, harvested, and processed through a bead-based pull-down on the reconstituted reporter. Annealing, BstXT reverse transcription at 65 C for 60 min, and the first Q5 PCR were all performed with the complex retained on beads, and the product was assessed by gel and verified by Sanger sequencing. The chimera migrated at the expected size of approximately 964 base pairs, reflecting the full barcode, tethering and handle cassettes of the cellular constructs (versus the short in vitro oligonucleotides), with the annealing junction near its center flanked by the 5x P22BoxB and 4x TAR arrays and the two barcodes.

**Figure 3.**
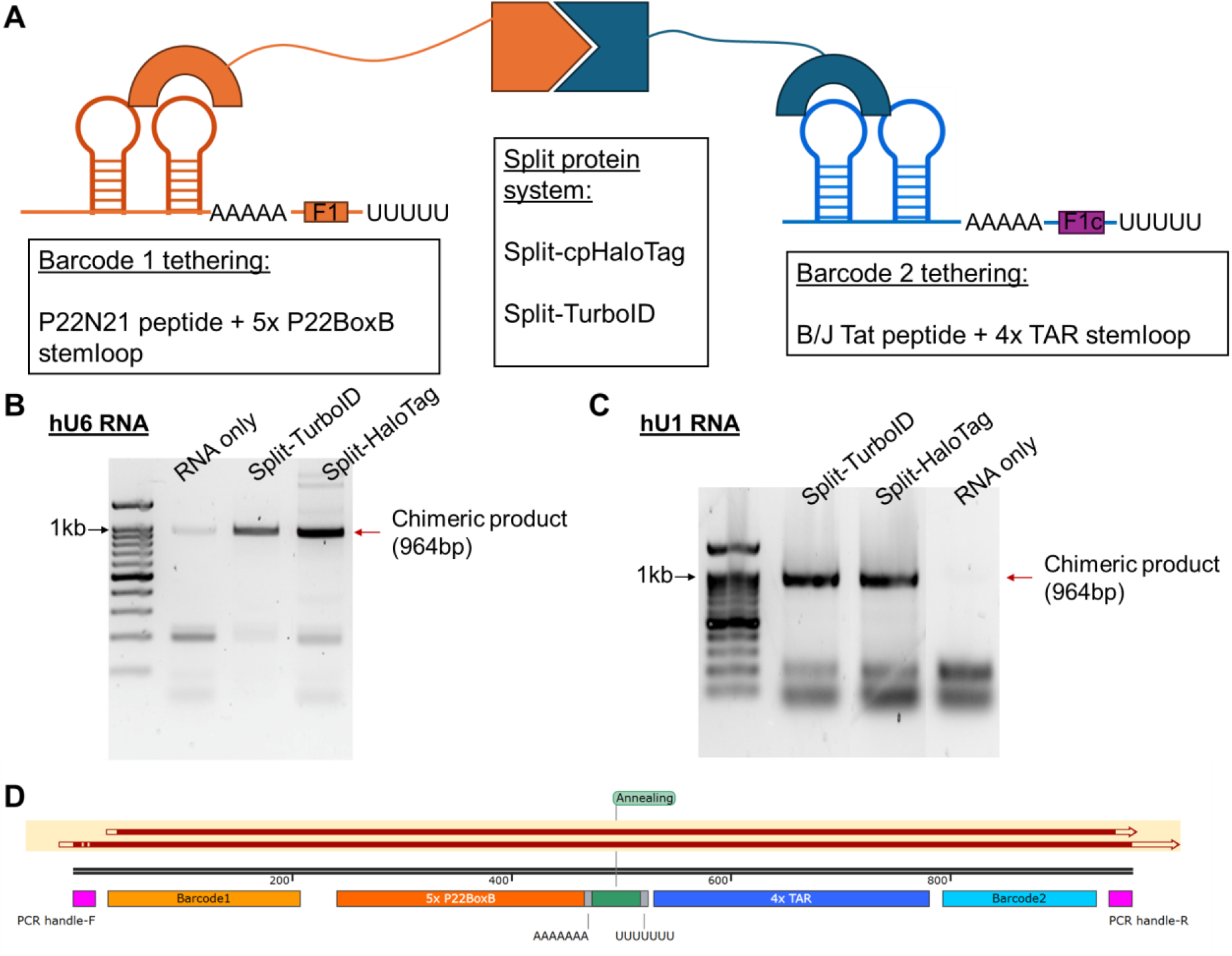
PAIR-link reads proximity of cellularly transcribed RNAs. *(A)* Reporter design. Each barcode RNA is tethered to one fragment of a split protein (split-cpHaloTag or split-TurboID) through the P22BoxB/P22N21-peptide and TAR/Tat modules; reconstitution co-localizes the RNAs and provides the pull-down handle. *(B)* Nuclear system (NLS-tagged split protein, U6-driven barcode RNA): RNA-only, split-TurboID and split-cpHaloTag conditions, resolved by gel; the chimera is approximately 964 base pairs. *(C)* Cytoplasmic system (NES-tagged split protein, U1-driven barcode RNA), same conditions. RNA-only gives no product in both systems. *(D)* Architecture of the 964 base-pair chimera (PCR handle, barcode 1, 5x P22BoxB, annealing junction, 4x TAR, barcode 2, PCR handle) and a representative Sanger trace confirming correct chimera are detected in the respective experiments.

We validated the nuclear and cytoplasmic reporters independently before attempting to run them together. A nuclear system, using NLS-tagged split protein with a U6-driven barcode RNA, and a cytoplasmic system, using NES-tagged split protein with a U1-driven barcode RNA, each produced the 964 base-pair chimera (Figure 3B, C). Both split reporters worked: split-TurboID recovered on streptavidin and split-cpHaloTag recovered on HaloTag beads each gave the product, while an RNA-only condition lacking the proteins gave none, as expected for a proximity-dependent reaction. We noted that recovery was most reproducible from freshly harvested material or material stored briefly at -80 C, which we treat as a controlled variable in current experiments.

### Toward multiplexed PAIR-link in distinct subcellular compartments

A reporter that operates in one compartment at a time is useful, but the larger goal is to read proximity in two compartments at once from a single cell population, so that nuclear and cytoplasmic events can be compared within one experiment. To test this design, we ran an eight-component experiment: a nuclear system (NLS-tagged split-TurboID with a U6-driven barcode pair) and a cytoplasmic system (NES-tagged split-cpHaloTag with a U1-driven barcode pair) were co-transfected, and products were recovered from total lysate or after nuclear/cytoplasmic fractionation (Figure 4A).

**Figure 4.**
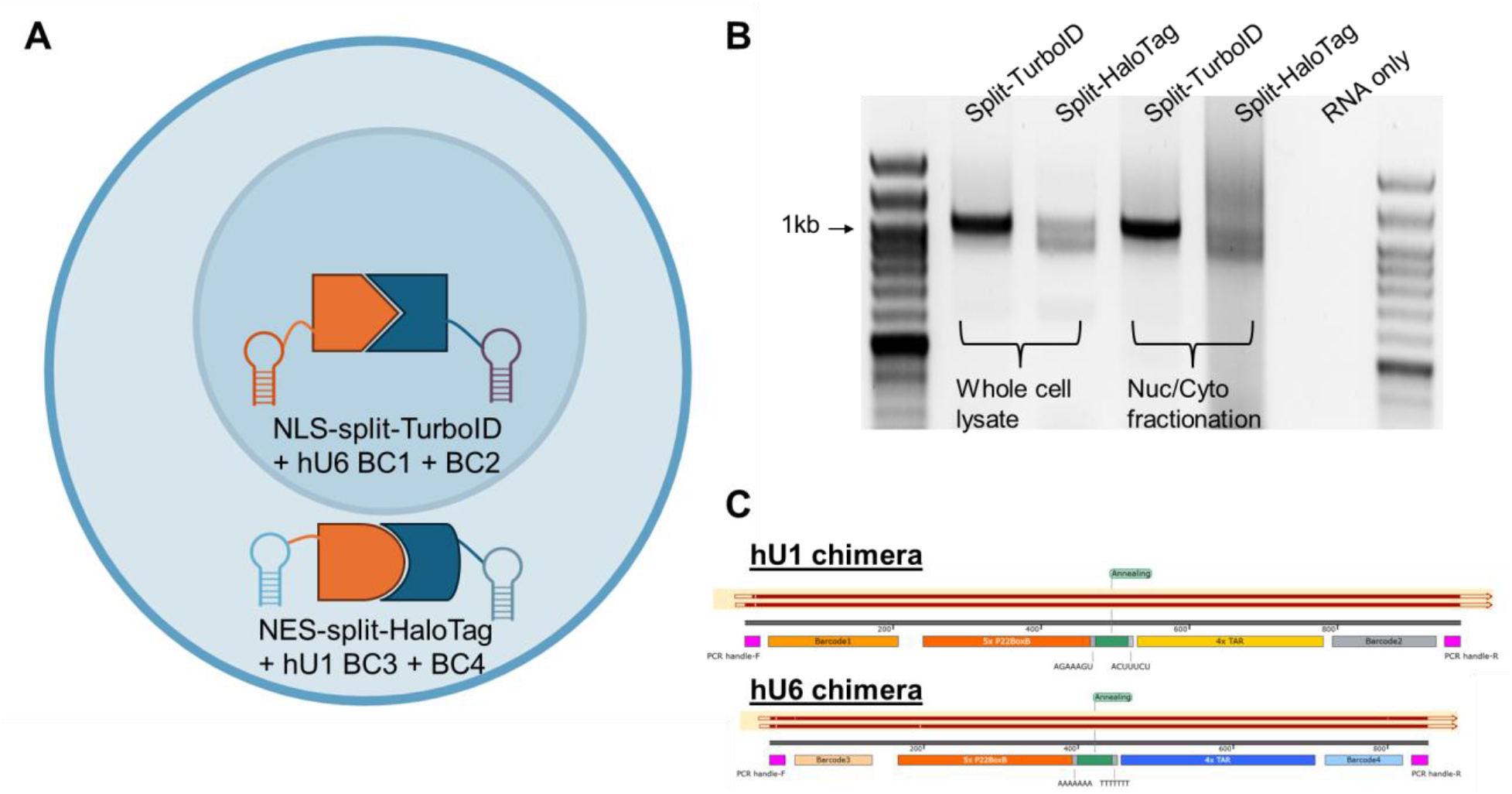
Toward multiplexed PAIR-link in distinct subcellular compartments. *(A)* Eight-component experiment: a nuclear system (NLS-tagged split-TurboID, U6-driven barcode pair) and a cytoplasmic system (NES-tagged split-cpHaloTag, U1-driven barcode pair) co-transfected in one cell population. The two systems use distinct annealing tags but share the BoxB and TAR tethering modules. *(B)* Products recovered from total lysate or after nuclear/cytoplasmic fractionation; the products of the two systems appear together rather than separated. *(C)* Representative Sanger traces confirming both types of chimeric products are produced.

Neither approach cleanly separated the products of the two compartments. Total lysate yielded both products together, and fractionation did not resolve them cleanly (Figure 4B, C). The cause is most likely due to molecular mingling rather than imperfect physical separation. We had given the two systems distinct annealing tags, so cross-talk at the RNA-RNA annealing step is excluded by design. What the two systems still share is the protein tethering: both use the same P22BoxB and TAR modules. A split-protein fragment from one system can therefore bind the barcode RNA of the other wherever the systems meet, which they do in total lysate and wherever the two compartments are not perfectly resolved. Because the shared tethers make the systems interchangeable at the protein-RNA binding step, no fractionation can fully prevent the cross-products, and this defines the design priority for the next stage of the method.

## Discussion

PAIR-link introduces a different way to turn proximity into sequence. By giving two RNAs complementary 3’ ends and letting an isothermal reverse transcriptase copy across the annealed junction, it fuses two separately transcribed molecules into a single cDNA in which the sequence of both partners is preserved, with no crosslinking, no fragmentation and no ligation. To our knowledge this is the first demonstration of such sequence-preserving linkage of two distinct, separately produced RNAs. The contrast with the established RNA-RNA interaction methods is the central point: those approaches recover partners as short, ligated fragments and pay an efficiency and bias penalty at the ligation step, whereas PAIR-link yields a contiguous product that is read directly.

Coupling the chemistry to a reconstituted RNP extends its use from RNA-RNA proximity to protein proximity. Because the two RNAs are tethered to two proteins, the chimeric product reports that the proteins were close, and the localization tags let the same reporter operate in defined compartments. The present study validates this readout when the protein modules are co-targeted to the nucleus or to the cytoplasm, which shows that the reporter functions across subcellular environments. Converting this into a quantitative protein-protein interaction assay, with interaction-dependent and interaction-independent controls, is the natural next step and is the application we consider most immediately useful.

Reading two compartments at once exposed the design feature that most limits the present system. Because both reporters use the same boxB and TAR tethering modules, their proteins and RNAs are interchangeable at the binding step, and neither total lysate nor fractionation can keep their products apart. The remedy is orthogonality at the tether, not better physical separation. The P22BoxB/P22N21-peptide and TAR/Tat pairs are two of several orthogonal viral RNA-peptide systems, and the bacteriophage MS2 and PP7 coat-protein/hairpin pairs are well established as a mutually orthogonal set usable in the same cell (Hocine et al. 2013). By rebuilding the second system from MS2 and PP7 in place of BoxB and TAR, the two systems become molecularly insulated, so that a fragment from one system cannot engage the barcode RNA of the other even within the same compartment. With orthogonal tethers in place, multiplexed and compartment-resolved PAIR-link should follow directly from the chemistry validated here, and because further orthogonal pairs are available the principle scales beyond two systems.

With orthogonal tethers in hand, the same architecture supports the applications we intend to pursue. Attaching a distinct barcode RNA to each protein of interest turns PAIR-link into a direct reporter of protein-protein interaction, where the ratio of chimeric to single barcodes provides a quantitative measure of how often two proteins meet, and where the availability of more than one orthogonal tether removes the restriction to pairs and opens a route to higher-order proximity among several proteins at once. Tethering one barcode to a protein and supplying the partner as an RNA would let PAIR-link verify protein-RNA interactions. The defined-3’-end requirement, which today is met by driving barcodes from Pol III promoters, points to native Pol III transcripts as the most tractable endogenous substrates: a small modification that presents a matching tag at their defined 3’ end would let PAIR-link record protein contacts on a Pol III RNA, or the contact between two Pol III RNAs, as a single sequence-preserving chimera. Extending the method to polyadenylated Pol II transcripts, which lack a defined end, is the harder and more general goal for our future research. We present the validated chemistry and cellular reporter here as the foundation for these directions.

## Materials and Methods

### Constructs

Barcode RNAs were driven from the human U6 or U1 promoters so that transcripts terminated at a defined 3’ end, with the F1 or F1c annealing tag placed immediately upstream of the terminator. One barcode RNA carried five copies of the P22BoxB hairpin and the other four copies of the TAR element. The cognate phage P22N21 peptide and B/J Tat peptide were each fused to one fragment of a split reporter protein, either split-cpHaloTag (cpHaloDelta paired with an Hpep peptide) or split-TurboID (TurboN and TurboC fragments), and tagged with an SV40 NLS for nuclear targeting or an NES for cytoplasmic targeting. Representative protein constructs included pCMV-SV40-NLS-cpHaloDelta-1xboxB-peptide and pCMV-SV40-NLS-Hpep1-3xTat, with the corresponding TurboN and TurboC pair for the split-TurboID reporter and the NES-tagged counterparts for the cytoplasmic systems. Barcode RNA constructs followed the form hU6- or hU1-[barcode]-5xP22BoxB-[F1 region]-4xTAR-[barcode], with short-end and long-end U1 3’ box variants tested.

### In vitro linking reactions

Synthetic barcode RNA oligonucleotides with complementary 3’ tails were annealed by controlled-temperature incubation. For the Bst3.0 reaction, reverse transcription was performed across a temperature and time series, followed by PCR with junction-flanking handle primers; the approximately 96 base-pair product was cloned into a landing plasmid, transformed into competent bacteria, and colonies sent for Sanger sequencing. For the BstXT reaction, reverse transcription was performed at 60 C and 65 C for 30, 40 and 50 min, amplified with Q5 polymerase and handle primers, gel-extracted, and similarly cloned and submitted for Sanger sequencing.

### Cell culture, transfection, and bead-intact pull-down

HEK293T cells were transfected with the barcode RNA and split-protein constructs. Cells were harvested and lysed, and the reconstituted reporter was captured by bead-based pull-down, on HaloTag resin for split-cpHaloTag or on streptavidin beads for biotinylated split-TurboID. Annealing, BstXT reverse transcription at 65 C for 60 min, and the first Q5 PCR were performed with the beads retained, to preserve the complex through processing. Products were resolved by agarose gel electrophoresis and verified by Sanger sequencing. An RNA-only condition lacking the protein constructs served as a negative control. Reactions used freshly harvested lysate or lysate stored briefly at -80 C.

### Subcellular fractionation

Two fractionation schemes were used. In the rapid scheme, cells were resuspended in polysome lysis buffer (PLB) on ice, centrifuged at low speed to collect the cytoplasmic supernatant, washed in NT-2 buffer, and the nuclear pellet was lysed in PLB; both fractions were cleared at 20,000 x g before immunoprecipitation. In the stringent scheme, cells were resuspended in hypotonic lysis buffer (HLB) on ice and centrifuged to collect the cytoplasmic fraction, the pellet was washed three times in HLB, and nuclei were lysed in nuclear lysis buffer (NLB); for one experiment half of the material was lysed in PLB in parallel to compare nuclear lysis conditions, and both fractions were cleared at 20,000 x g before immunoprecipitation. Buffer compositions are given in Table 1.

**Table 1.** Buffer compositions.

| Buffer | Composition |
| --- | --- |
| 10x PLB stock | 1 M KCl, 50 mM MgCl<sub>2</sub>, 100 mM HEPES-NaOH pH 7, 5% NP-40 |
| PLB working | 1x PLB, 1 mM DTT, RNaseOUT (80 U/mL), 1x protease inhibitor |
| 5x NT-2 | 250 mM Tris-HCl pH 7.4, 750 mM NaCl, 5 mM MgCl<sub>2</sub>, 0.25% NP-40 |
| NET-2 (from 1x NT-2) | 1x NT-2, 20 mM EDTA pH 8, 1 mM DTT, 1:200 RNase inhibitor |
| HLB | 10 mM Tris-HCl pH 7.5, 10 mM NaCl, 3 mM MgCl<sub>2</sub>, 0.3% NP-40, 10% glycerol, 1:200 RNase inhibitor, 1x protease inhibitor |
| NLB | 20 mM Tris-HCl pH 7.5, 150 mM KCl, 3 mM MgCl<sub>2</sub>, 0.3% NP-40, 10% glycerol, 1:200 RNase inhibitor, 1x protease inhibitor |

## Notes

### Competing Interest Statement

The authors have declared no competing interest.

